# The PEDtracker: an automatic staging approach for *Drosophila melanogaster* larvae

**DOI:** 10.1101/2020.09.30.320200

**Authors:** Isabell Schumann, Tilman Triphan

## Abstract

The post-embryonal development of arthropod species, including crustaceans and insects, is characterized by ecdysis or molting. This process defines growth stages and is controlled by a conserved neuroendocrine system. Each molting event is divided in several critical time points, such as pre-molt, molt and post-molt, and leaves the animals in a temporarily highly vulnerable state while their cuticle is re-hardening. The molting events occur in an immediate ecdysis sequence within a specific time window during the development. Each sub-stage takes only a short amount of time, which is generally in the order of minutes. To find these relatively short behavioral events, one needs to follow the entire post-embryonal development over several days. As the manual detection of the ecdysis sequence is time consuming and error prone, we designed a monitoring system to facilitate the continuous observation of the post-embryonal development of the fruit fly *Drosophila melanogaster*. Under constant environmental conditions we are able to observe the life cycle from the embryonic state to the adult, which takes about ten days in this species. Specific processing algorithms developed and implemented in Fiji and R allow us to determine unique behavioral events on an individual level – including egg hatching, ecdysis and pupation. In addition, we measured growth rates and activity patterns for individual larvae. Our newly created RPackage PEDtracker can predict critical developmental events and thus offers the possibility to perform automated screens that identify changes in various aspects of larval development. In conclusion, the PEDtracker system presented in this study represents the basis for automated real-time staging and analysis not only for the arthropod development.

## Introduction

Ecdysis or molting is the most important feature of ecdysozoan species including arthropods, nematodes and other relatives (Telford et al., 2008). The whole body surface of these animals is surrounded by a chitinous cuticle which hardened, for example, to an exoskeleton in the whole group of arthropods (Hadley, 1986; Ewer, 2005). Therefore, for a successful growth from juveniles to adults the animals have to shed and renew their body surface throughout their post-embryonal development (Nijhout, 2013). The renewing process is controlled by a highly conserved neuroendocrine system which has been well described in several species, especially in arthropods such as the crab *Portunus trituberculatus,* the butterfly *Manduca sexta* and the fly *Drosophila melanogaster* (Xie et al., 2016, Riddiford et al., 1999; Rewitz et al., 2007;). Beside the renewing of the cuticle, some insect groups further evolved a total re-organization (metamorphosis) from the last larval stage to the adult during a pupal stage (holometabolism). Consequently, the post-embryonal development of holometabolous insects is characterized by both, molting and metamorphosis (Weeks and Truman, 1984; Truman, 1996).

Although the molecular pathway and regulation of molting-related molecules are well understood, the question arises whether different factors such as the physiological state of the animals, food choice and environmental stimuli influence the post-embryonal development and directly affect the larval endocrine system and growth (Koyama et al., 2014). While hormones such as 20-hydroxyecdysone, juvenile hormone and insulin are well known to function in molting and growth especially in insects (Chang, 1985; Riddiford et al., 2003; Beckstead et al., 2005; Lin and Smagghe, 2019), the role and importance of the critical weight especially for the initiation of metamorphosis is still not fully understood (Robertson, 1963; Nijhout and Williams, 1974; Davidowitz et al., 2003; Mirth et al., 2005).

To get a better understanding of the physiological state, molecular background and motivation of the individual animal during specific developmental time points, such as molting or metamorphosis, staging of juveniles or larvae represents an important approach. For example, larvae of holometabolous insects can be manually classified into developmental stages by morphological characteristics such as the differentiation of the mouth hooks or anterior and posterior spiracles (according to, e.g. Okada, 1963; Schubiger et al., 1998; Vaufrey et al., 2018). Another approach for staging juveniles or larvae is to allow females to lay eggs for some hours on a food media and examine larvae after a set time interval, for example every 12 hours (according to, e.g. Burmester et al., 1999). However, both manual detection approaches are time-consuming. In addition, specific larval stages and developmental time points are difficult to catch as the main ecdysis behavioral sequence takes only a comparatively short time window of a few minutes. The molting cycle in general comprises four consecutive phases: (1) intermolt; the time between two molting events (2) pre-molt; the time-window just before the main molt (3) molt; the time-window where larvae shed their cuticle and (4) post-molt; the time-window right after the main molt (Locke, 1970). In the fly *D. melanogaster* the ecdysis behavioral sequence as part of the molting cycle is described to last only 30 minutes and is divided in four parts – the occurrence of the double mouth hooks and vertical plates, renewing of the tracheal system, pre-ecdysis and ecdysis (Park et al., 2002).

Our long-term goal is the investigation of individual molecular and behavioral changes during the molting cycle of the animal. As the main ecdysis takes about five minutes of a single larval stage of *D. melanogaster,* the first aim was to find a tool for precisely timing these stages for investigations on development-associated processes such as sensation and food choice. Using *D. melanogaster,* we established a monitoring system throughout the whole post-embryonal development – from egg to pupa – over ten days (Dewhurst et al., 1970a). Under constant environmental conditions (25°C and 65% humidity) and a yeast-sugar diet, we observed the life cycle from egg to pupa with a camera and a framerate of 3 frames/minute. We subsequently analyzed the videos with the focus on the molting cycle throughout the post-embryonal development such as the main molting sequence on an individual level. Our results reveal insights into larval behavior regarding activity and growth. We clearly observe the specific activity pattern during the molting cycle and the individual growth in size from young to old larval stages. Consisting of a video recording set up and newly developed analysis scripts (in Fiji and R), the PEDtracker (= post-embryonal development tracker) forms the basis for a future real-time tracking system for the prediction of developmental stages which could also be used for various other insects and their relatives.

## Material and Methods

### Animal husbandry and fly strains

All experiments were performed with the wild-type *Drosophila melanogaster* strain *Canton-S*. Flies were kept on standardized cornmeal medium at 25°C and 65% humidity under a 14:10 light:dark cycle. Adult flies were transferred to new food vials every 72h.

### Egg laying, tracker preparation and data recording

For egg laying small Petri dishes (4 cm diameter) were filled with a 3% agarose (VWR life science; type number: 97062-250), 3% sucrose (Merck KGaA; type number: 107687), and 30% apple juice (Edeka) mix. To entice female flies to oviposition one drop of yeast was put on top of the plate. Flies were allowed to lay eggs for at least two hours and then, one egg per chamber was transferred to the larval bed. The bed was prepared as previously described (Szuperak et al., 2018) with the SYLGARD^®^ 184 Silicone Elastomer Kit (type number: 24001673921) and a size of 8.3 cm in length and 5.7 cm in depth in a self-made 3D-printed template (Renkforce 3D-printer RF1000; material PLA). Each bed contains of 24 chambers (4×6) with each 1 cm in diameter. Before placing eggs, each chamber was filled with 100μl food medium containing a mix of 2% agarose (VWR life science; type number: 97062-250), 2% sucrose (Merck KGaA; type number: 107687), and three drops of fresh yeast. To avoid mold a mixture of 0.1% methylparaben-ethanol (methylparaben: Carl Roth GmbH + Co. KG type number: 3646.4; ethanol: CHEMSOLUTE^®^ type number: 2273.1000) was added to the food medium. To avoid an escape of larvae, the bed was covered with clear film and two layers of glass plates. The bed was then placed in a custom-built climate chamber (workshop of the University of Konstanz) on a glass table with constant LED-light (KYG light-table) from below (Fig. 1). The climate chamber provided a constant temperature (23–25°C) and humidity (60–65%) for our experiments. Pictures were taken every 20 seconds over a period of at least ten days with a 25mm lens using a Basler camera (acA2040-25gm; type number: 105715) with a resolution of 2048×2048.

**Figure 1.**
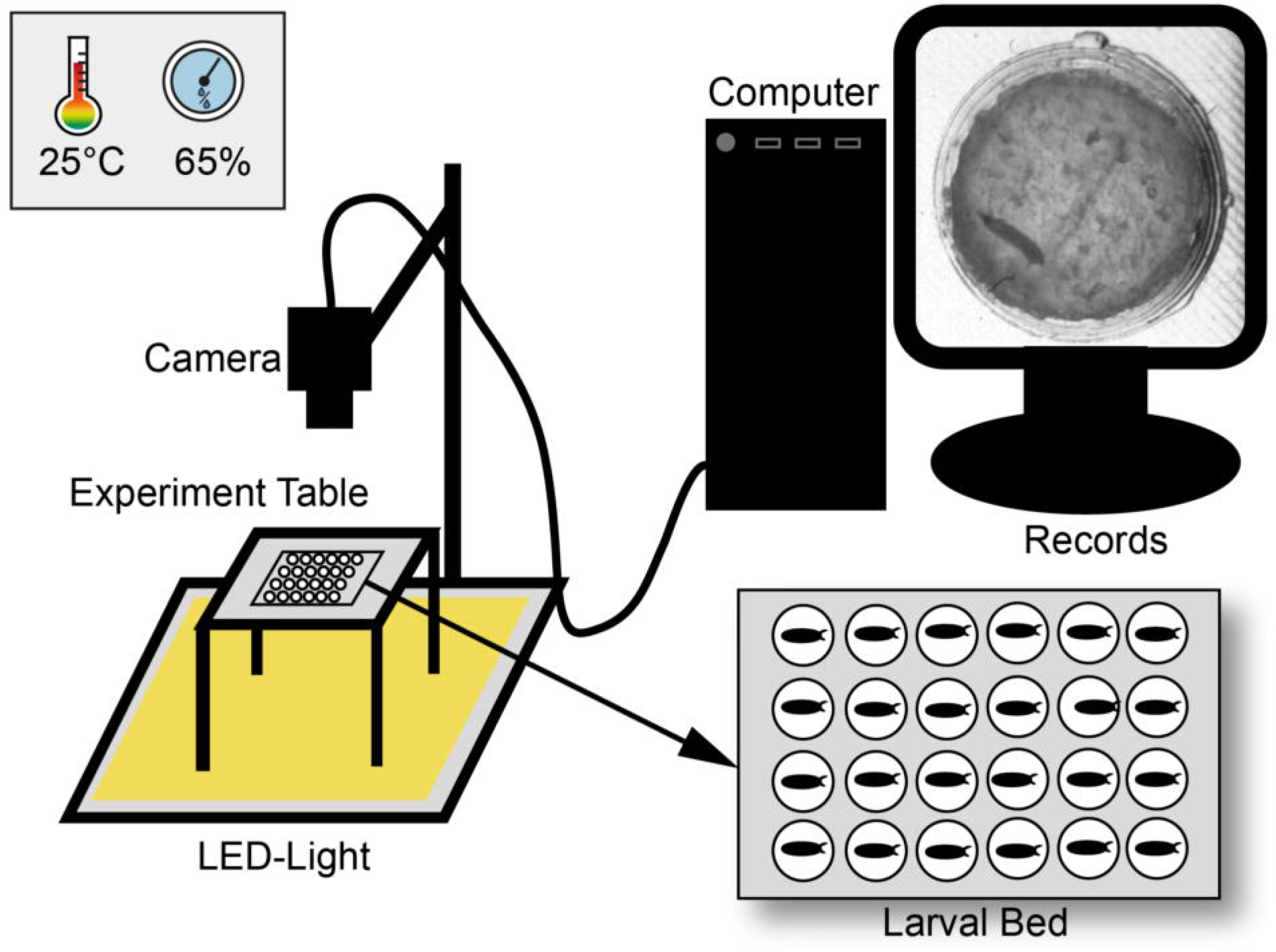
Scheme of the experimental set-up. Larvae were observed over ten days under constant light in a climate chamber with constant abiotic conditions. For observation three pictures per minute were recorded with a camera and saved with a custom-made software. Pictures were then combined to a video sequence and analyzed with custom-made scripts in ImageJ and R.

### Data analysis

For data analysis pictures were processed as individual six-hour video sequences (1080 frames) with a custom-made Fiji script (Schindelin et al., 2012; see Supplementary Material S3). We first defined regions of interest (ROIs) marking the individual chambers. For each ROI a median background image was created and subtracted from the cropped video. The resulting images were thresholded and then analyzed with specific measurement settings (Tab.1) in Fiji.

**Table 1.** Measurements used in ImageJ for video analysis (full description via https://imagej.nih.gov/ij/docs/guide/146-30.html#toc-Subsection-30.7).

| Measurements | Description |
| --- | --- |
| **Shape Parameters** |  |
| *Aspect ratio* | $\frac{Major\ Axis}{Minor\ Axis}$ |
| *Circularity* | $4\pi * \frac{Area}{Perimeter^2}$<br>0 = no circle<br>1 = circle |
| *Roundness* | $4 * \frac{Area}{\pi * Major\ Axis^2}$ |
| *Solidity* | $\frac{Area}{Convex\ area}$ |
| **Other parameters** |  |
| *Area* | Selection in square pixels |
| *Centroid* | Center of point in the selection |
| *Fit ellipse* | Add ellipse to the selection by calculating primary and secondary axis |

After image processing in Fiji (Fig. 2), we followed up with error correction and further in-depth analysis in a custom-written R script using R version 3.6.1 (R Core Team, 2019) in RStudio (R Studio Team, 2019; see Supplementary Material S4). For each larval stage, parameters for size and shape of valid objects were defined (Tab. 2, Fig. 3). To determine the length of each larva the major axis of each object was scaled to the diameter of each chamber (~ 300 px = 10 mm). Growth rates were then calculated as follows:

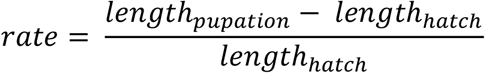

**Figure 2.**
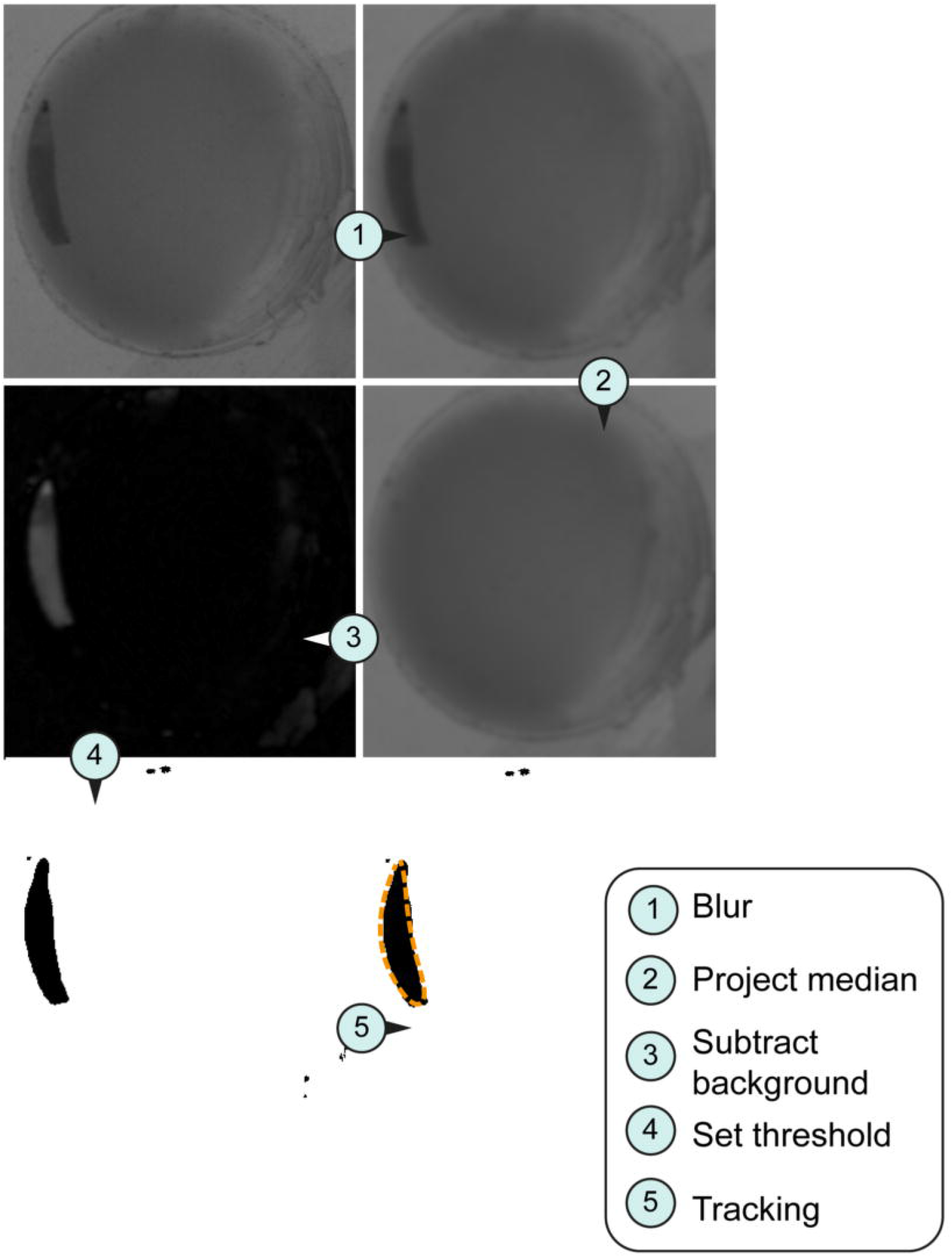
Image processing for tracking system. Larval stage three were used all imaged pictures. (A) Original section of a video sequence. Note individual larva in the chamber. (B) Section of a video sequence. Picture were smoothed using convolution with Gaussian function. (C) Median calculation of an image shows the median intensity over all images in a stack. (D) Implementation of an arithmetic and logical operation. We used the difference between the source (img1) and destination image (img2) – imgX = | img1 – img2 |. (E) Set threshold dependent on the larval stage (L1, 10; L2, 20; L3, 30). (F) Orange dashed line represents the silhouette of the object of interest in the section.

**Figure 3.**
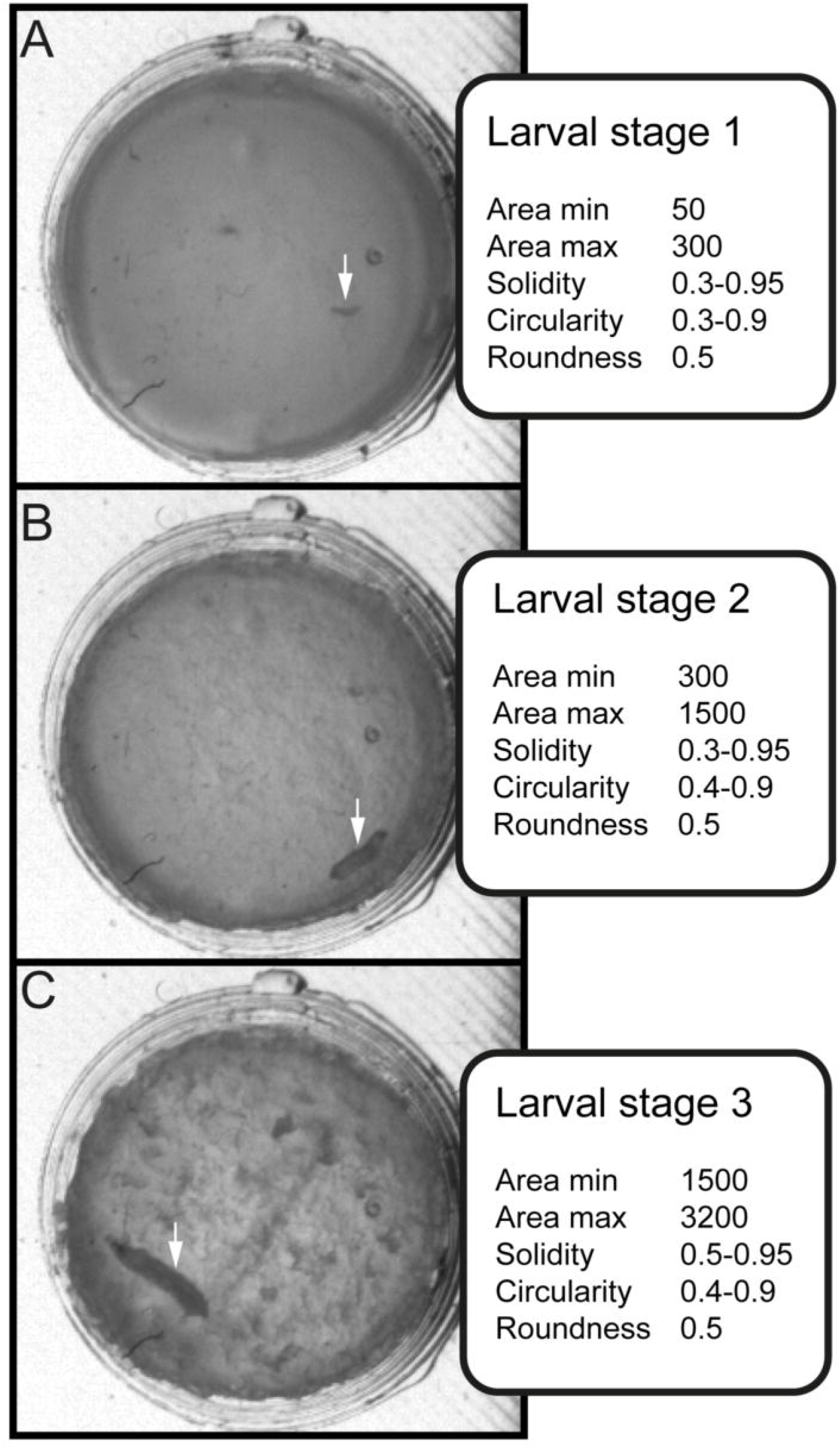
Stage-specific parameters for data sorting and evaluation. Three values for shape description – solidity, circularity and roundness – are used in combination with the area for differentiate between larval stages. (A) Arrow indicate position of larval stage one. Surface of individual larva ranges from 50 to 300 square pixels. (B) Arrow indicate position of larval stage two. Surface of individual larva ranges from 300 to 1500 square pixels. (C) Arrow indicate position of larval stage three. Surface of individual larva ranges from 1500 to 3000 square pixels.

**Table 2.** Comparison of developmental time points. Abbreviations: AEL, after egg laying; AH, after hatching

|  | **PEDtracker** | **Literature** |
| --- | --- | --- |
| **Hatch** | 0.7 days AEL | ~ 0.8 days AEL<br>(see Markow et al., 2009) |
| **1<sup>st</sup> molt** | 1.9 days AH | ~ 1 day AH<br>(see Parkin and Burnet, 1986) |
| **2<sup>nd</sup> molt** | 3.5 days AH | ~ 2 days AH<br>(see Parkin and Burnet, 1986) |
| **pupation** | 7.8 days AH | ~ 3.5 days AH<br>(see Parkin and Burnet, 1986) |
| **period of larval development (hatch–pupa)** | 7.8 days | ~ 4 days<br>(see Powsner, 1935) |

To determine larval movement, the Euclidian distance between the centroids (X_1_, X_2_; Y_1_, Y_2_) in two successive frames was calculated through Pythagorean function

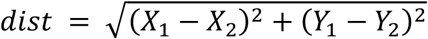

Next, the relative movement over time was used to get an activity pattern for the individual larvae. Data frames containing analyzed particles were then sorted, merged and plotted. We show examples of the molting behavior of two larval stages in time lapse videos (recorded at 3 fpm; played back at 3 fps; see Supplementary Material S1 and S2). Final figures were edited with Adobe Illustrator CS5 (San Jose, CA, USA).

## Results & Discussion

As a basis for our tracking setup we used the LarvaLodge (Szuperak et al., 2018) which itself is based on earlier work in *Caenorhabditis elegans* (Churgin et al., 2017). In their experiment up to 20 larvae can be monitored simultaneously for several hours. For our even longer approach – monitoring the whole post-embryonal development of *D. melanogaster* over the course of several days – we had to overcome multiple challenges. Our first task was to improve the setup to enable proper larval develop over several days. For this we had to address humidity issues and prevent the growth of mold. We settled on a frame rate of three frames per minute as a middle ground to still be able to measure activity but also record for a timespan of more than a week without generating a vast amount of data. One typical experiment contains about 60,480 images (14 days; lossless compression as .png-images). The first step of processing was performed in ImageJ where the target objects (i.e. the larvae) are extracted from the video. Here we had to address the problem of the dramatic change in size of the larvae over the course of their development as well as the change in the environment over several days. Difficulties occurred with food drying out or larvae digging into the food. Further improvement of the quality of our data was then performed in the secondary processing implemented with R in RStudio.

Our results reveal a mean hatching time point of 18 hours after egg laying and a nearly similar size of about 0.5 millimeter for young L1 *Canton-S* larvae (Fig. 4, 5). Interestingly, the later the developmental stage the higher is the variability of the time point of molting events and the respective body sizes of larvae (Fig. 4A, C; Fig. 5). Our results indicate a mean lifespan of *D. melanogaster Canton-S* larvae of 7.8 days from egg hatching to pupation (Fig. 4A; Fig. 7). This result is in contrast to previous studies which revealed that *D. melanogaster* larvae went under pupation five days after egg laying (Tab. 2, Dewhurst et al., 1970b; Ashburner, 1989; Casas-Vila et al., 2017). Our results further indicate less stringency in developmental time points and life spans of single larval stages in comparison with the literature (Fig. 7). Whereas the mean hatching time after egg laying is in line with previous studies (about 18 hours; 0.7 days; Markow et al., 2009), our results indicate the first molting event 2.6 days after egg laying (= 1.9 days after hatching) and the second molting event 4.2 days after egg laying (= 3.5 days after hatching). Previous studies indicated that larvae of the same age not pupated at the same time (Casares and Carracedo, 1987). Thus, the life cycle of *D. melanogaster* might be variable to some degree and dependent on different lifehistory traits and not only on the expression of molting related molecules such as ecdysone and juvenile hormone.

**Figure 4.**
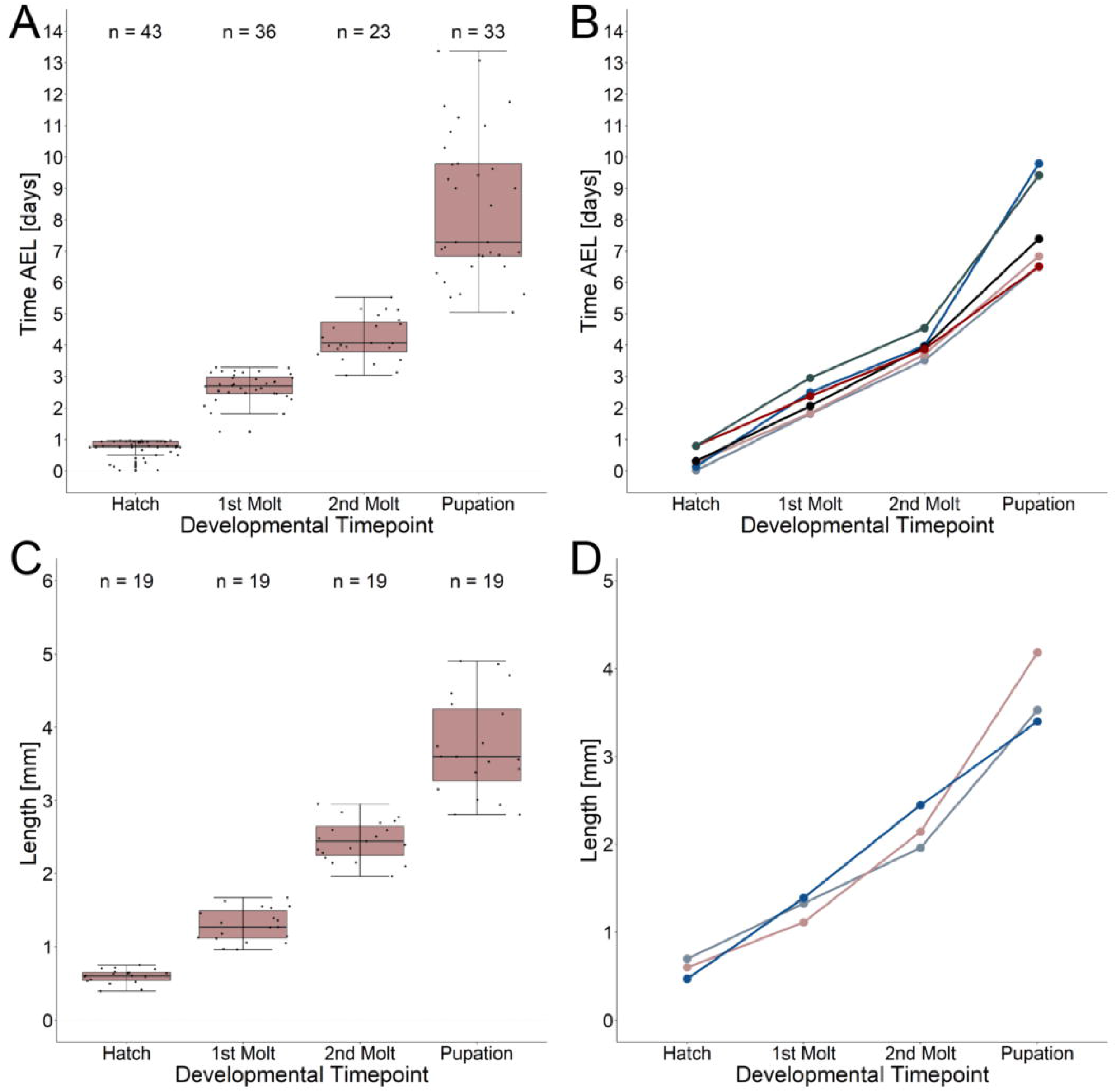
Developmental time points and related growth in length. (A) Time after egg laying and corresponding developmental time points. Note the high variation (five to ten days after egg laying) of the pupation event. (B) Time after egg laying and corresponding developmental time points for five single larvae. Note the high variation of the pupation event independent on the hatching time point. (C) Growth in length from egg hatching to pupation and related developmental time points. Note the high variation of larval length at the pupation event. *Growth rate^median^* = 4.9. (D) Growth in length and related developmental time points for three single larvae. Note the highest growth in length in larval stage three. *Growth rate 1(blue line)* = 6.2; *growth rate 2 (light red line)* = 5.9; *growth rate 3 (grey line)* = 4.06.

**Figure 5.**
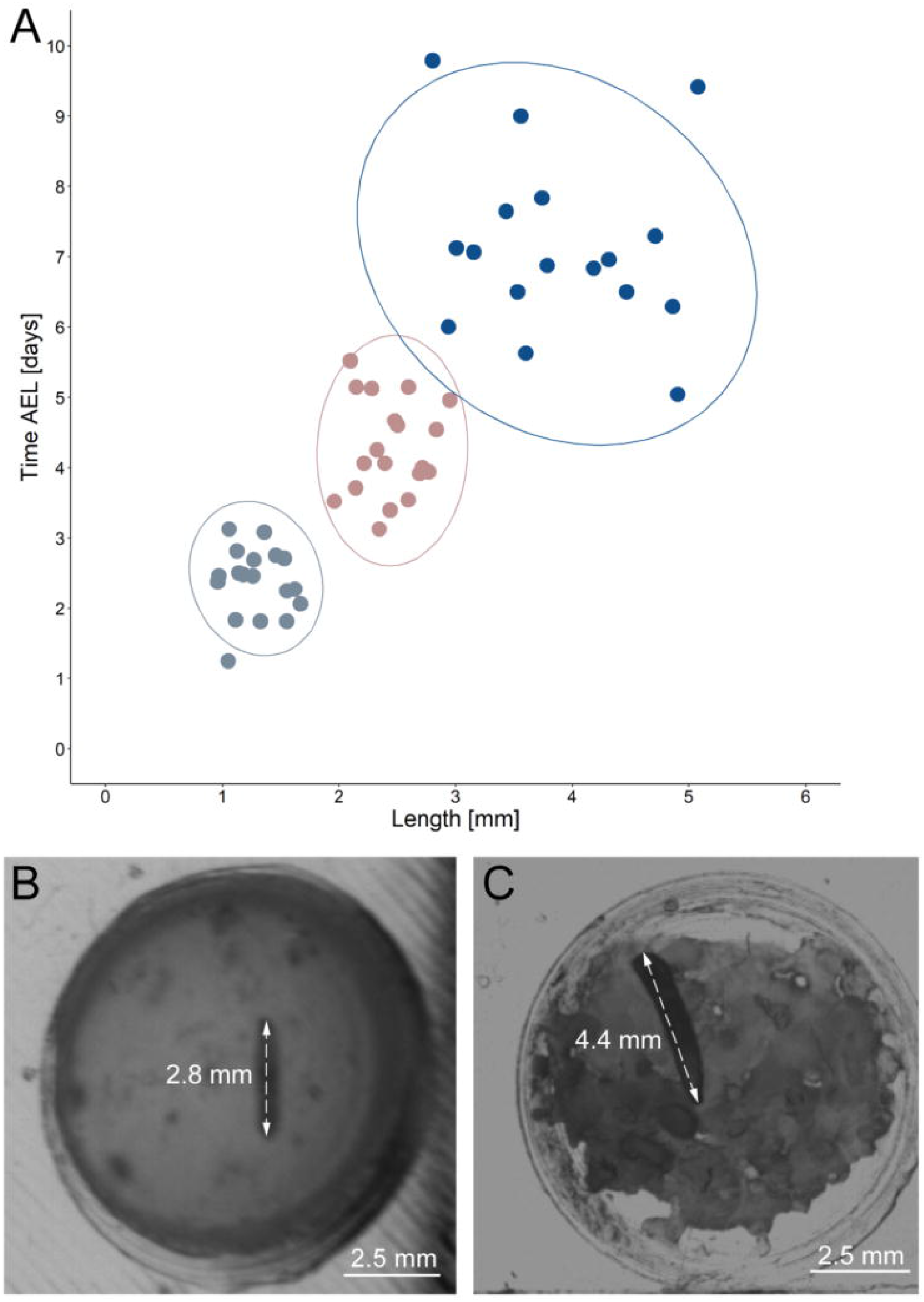
Correlation between developmental time and size. (A) Y-axis indicate time after egg laying in days; x-axis represent larval length in mm. Light grey dots indicate larval stage one, light red dots indicate larval stage two and blue dots indicate larval stage three. Ellipses represent 95% confidence level. (B) Arrows indicate the length of a small late stage three larvae, approximately one hour before pupation.. (C) Arrows indicate the length of a large late stage three larvae approximately one hour before pupation.. Abbreviations: AEL, after egg laying; mm, millimeter.

Previous studies showed that the developmental period of drosophilid species depends on environmental stimuli such as temperature, humidity or light. The developmental period successively decreases to seven days depending on the increase of the temperature up to 28 degree (Ashburner and Thompson Jr, 1978; Al-Saffar et al., 1996; Tochen et al., 2014). Moreover, the highest survival rate (88–97%) for pupae of *D. melanogaster* is described for 60 to 100% relative humidity (Elwyn et al., 1917; Al-Saffar et al., 1996). But then, the sensitivity of larvae to temperature and humidity differ between drosophilid species (Geisler, 1942; McKenzie and Parsons, 1974). In contrast to the effect of temperature and humidity, the role of light on the developmental period and larval circadian clock is still under debate. Some studies indicated that permanent night has no effect on the animals, but in other cases the culture in constant darkness revealed a reduced longevity (Payne, 1910, 1911; Erk and Samis, 1970). Similar effects have been shown for culture in constant light (Allemand et al., 1973).

Investigation of dietary influence on the developmental period in larvae further implicated that the food composition has a prevalent effect on the life span and body parameters of drosophilids (Anagnostou et al., 2010; Ormerod et al., 2017). The developmental period increases with the linear availability of carbohydrates (negative correlation) and decreases (positive correlation) with the linear availability of proteins. Consequently, larvae are capable to differentiate their needs of nutrients and regulate their dietary intake towards a minimal developmental period (Rodrigues et al., 2015). In two drosophilid species *D. melanogaster* and *D. subobscura* a correlation between the metabolic rate, the developmental period and social parameters was observed (Marinkovié et al., 1986; Cluster et al., 1987; Hoffmann and Parsons, 1989; Sevenster and Alphen, 1993). Additional investigation on larval life, mortality and pupal viability in *D. melanogaster* and *D. simulans* revealed a correlation between the density of larval numbers and the developmental period – high density and crowding of larvae in culture leads to a longer developmental period in both species (Powsner, 1935; Miller, 1964). To conclude, a decrease or increase of the developmental period of *D. melanogaster* is caused by different environmental stimuli and sociality. However, due to our results and experimental set-up, the examination of a single larva in one chamber on basic food medium with optimal temperature (25°C) and humidity (65%), we assume that the metabolic rate might influences the developmental time rather than social interactions or competition in their environment.

Our results further indicate a high variability of the size, especially for L3 *Canton-S* larvae. Whereas young L1 *Canton-S* larvae showed a similar size (about one millimeter) until the first molting event, late wandering L3 *Canton-S* larvae (shortly before pupation) differ in size from about three to five millimeters (Fig. 4C, Tab. 3). Therefore, we plotted larvae on an individual level to observe the time span and size between egg hatching and pupation on an individual level. Our results indicate that the pupation time point and the size of late L3 *Canton-S* larvae are independent of the egg hatching time and size (Fig. 4, 5). We further assume that the critical weight, or at least critical size, is more important for younger larvae than for older and therefore might play a key role for the initiation of molting (Fig. 5). Our correlation between the time after egg laying and larval size showed that also small L3 larvae of the *Canton-S* strain went under pupation after a relatively long developmental time (Fig. 5A). This result indicates that also smaller L3 larvae are able to pupate and therefore the size might not be the only initiator for pupation and metamorphosis (Fig. 5B, C). In comparison with other wild-type strains the variability of size in *Canton-S* larvae is also higher (Vaufrey et al., 2018).

**Table 3.** Comparison of larval body length. Abbreviations: L1, larval stage one; L2, larval stage two; L3, larval stage three.

|  | **PEDtracker**<br>**(larvae grew on basic medium)** | **Literature**<br>**(larvae grew on wild-type yeast)** |
| --- | --- | --- |
| **young L1** | 0.6 mm | ~ 1 mm<br>(see Parkin and Burnet, 1986) |
| **late L1 (1<sup>st</sup> molt)** | 1.3 mm | ~ 1.5 mm<br>(see Parkin and Burnet, 1986) |
| **late L2 (2<sup>nd</sup> molt)** | 2.5 mm | ~ 3 mm<br>(see Parkin and Burnet, 1986) |
| **late L3 (pupation)** | 3.8 mm | ~ 4.5 mm<br>(see Parkin and Burnet, 1986) |

The characterization of specific developmental time points and larval size are important parameters for the investigation of the body constitution as well as the behavior during the development. Looking at an example of larval activity over a period of about three hours we can see that the larva crawls through the chamber and stops at frame 213 (video position ~ 1:20 min, see Supplementary Material S1) and then rests in the same place for almost 30 minutes. Interestingly, after 14 minutes the larva turns to the side and rests there for another 15 minutes. During both resting periods, the record indicates alternating phases of resting, pulsation and contraction (frames 213–300, video position 1:20-1:40 min). After comparing the video to previous studies we suggest that the observed resting phase in the video sequence correspond to the ecdysis behavioral sequence of the *D. melanogaster* larva (Park et al., 2002). We further assume that the first resting phase corresponds to tracheal molt and air filling, and the second one with stronger pulsation and contractions to pre-molt and main molt. The same pattern occurs during the ecdysis behavioral sequence of L2 to L3 (see Supplementary Material S2). We analyzed both ecdysis behavioral sequences for larval activity (Fig. 6). The activity patterns for both larval stages reveal a decrease of movement during molting events. Whereas larvae move up to 5 mm in the intermolt phases, they slow down to at least 0.5 mm in the pre-molt and main molt stage (Fig. 6A–D).

**Figure 6.**
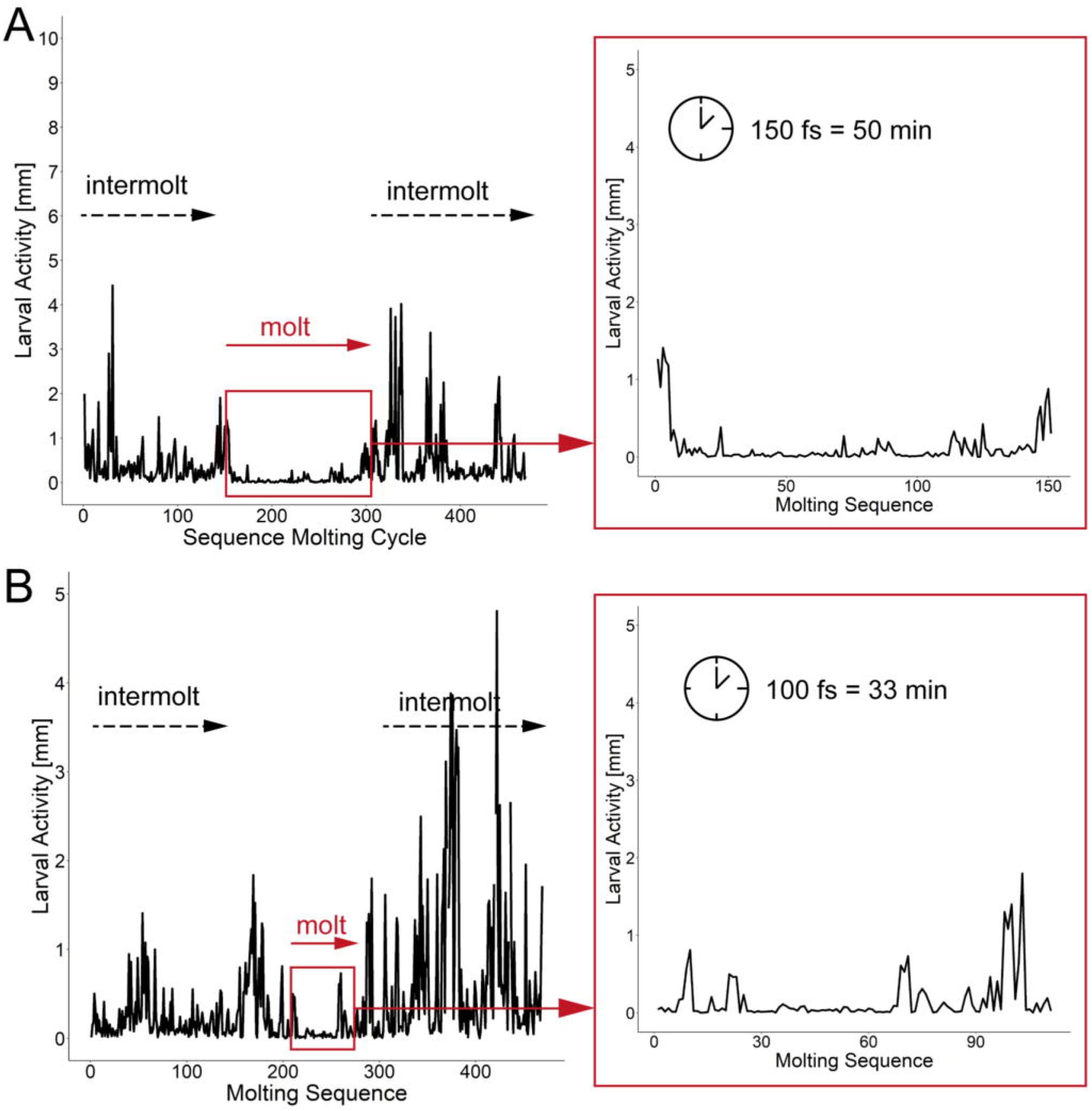
Activity patterns of two developmental stages and related molting events. (A) Activity pattern (in mm) and molting sequence of larval stage one. Insert represents the activity pattern (in mm) of larval stage one during the main molting event. Note a time window of 150 fs of low activity of the larva. (B) Activity pattern (in mm) and molting sequence of larval stage two. Insert represents the activity pattern (in mm) of larval stage two during the main molting event. Note a time window of 100 fs low activity of the larva. Abbreviations: fs, frames; min, minutes.

In this study we followed *D. melanogaster* larva during several stages of development from egg hatching to pupation over up to 14 days (Fig. 7). We defined tracking parameters to identify the larva most consistently in our environment and over a large range of body sizes. We found and analyzed critical developmental events like molting while looking at the activity and growth of the larva (Fig. 7). With the combination of a video-sequence and a particle analyzer we are able to manually detect important developmental stages which will be the basis for a future real-time tracking system. For future improvement of the PEDtracker system, the analyzed particles regarding larval size and activity we presented in this methodology illustrate the basis for a custom-made software program for the analysis of insect larvae and prediction of behavioral events. Our focus was on getting data for the future implementation of such a real-time tracking system with an integrated molting detector for the prediction of important development time points such as molting or metamorphosis.

**Figure 7.**
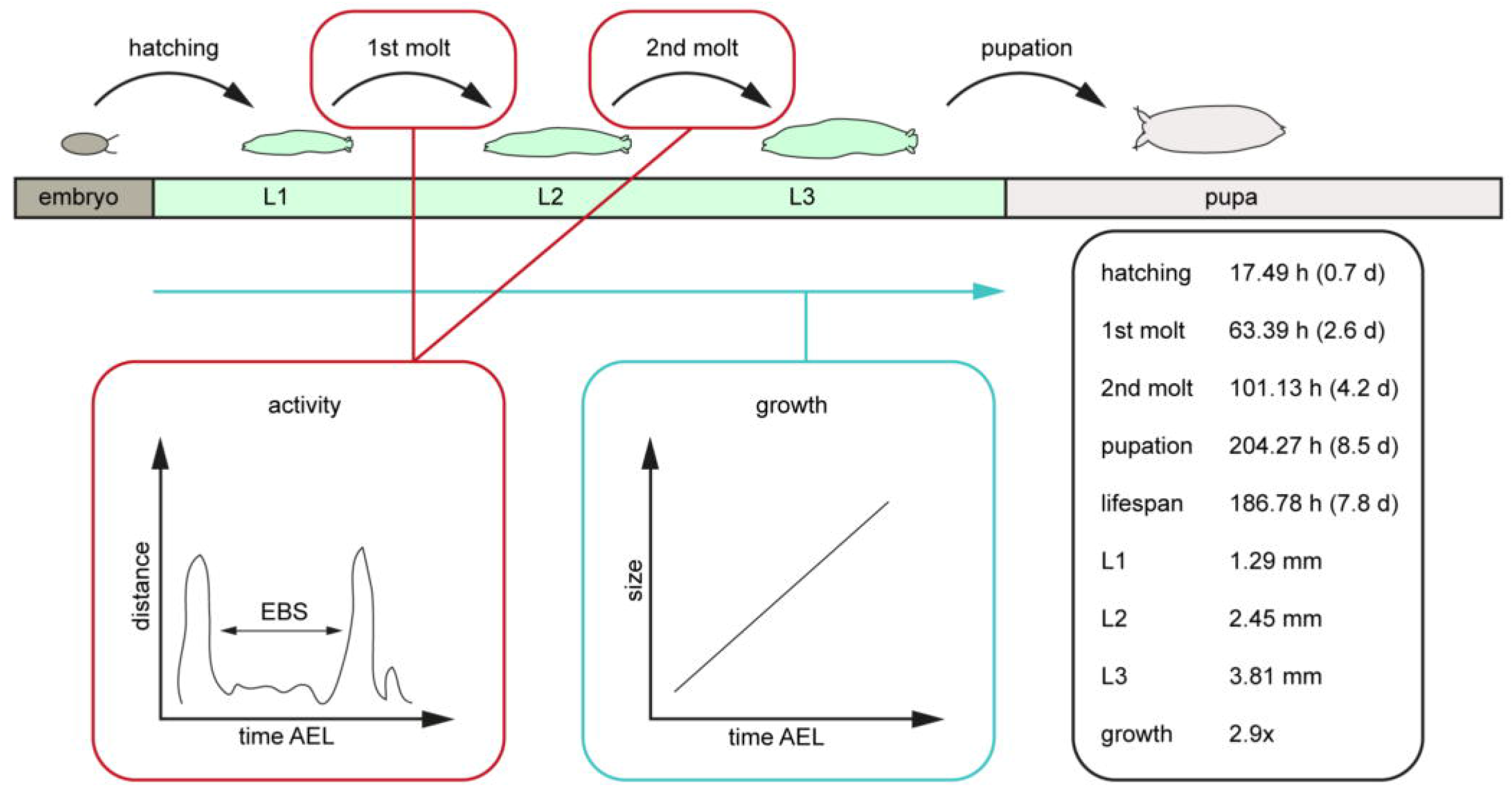
Scheme of the post-embryonal development of *D. melanogaster* and related values generated by the PEDtracker. The given values represent the average values of relevant developmental time points after egg laying such as the larval lifespan, the average length of all larval stages in length and the related growth rate. Abbreviations: AEL, after egg laying; d, days; EBS, ecdysis behavioral sequence; h, hours; L1, larval stage one; L2, larval stage two; L3, larval stage three; mm, millimeter.

Taken together, our PEDtracker system provides a novelty in tracking systems for the observation of the whole post-embryonal development on an individual level which is not only suitable for insects but also for other molting animals such as chelicerates, nematodes and other ecdysozoans. Besides the usage in the observation of developmental time points, the PEDtracker represents a useful tool for further molecular and behavioral experiment such as the culture of different genotypes under different food regimes.

## Data Availability Statement

Datasets generated for this study are available on request to the authors. Fiji script (PEDtrack.ijm) and R package are available on: https://github.com/IzzySchu/PEDtracker.

## Acknowledgements

We are thankful to Andreas S. Thum who supported the whole project and made helpful comments on the manuscript. We are furthermore grateful to the whole Department of Genetics for support and comments on the establishment of the experimental procedure. We are also thankful to Ingo Kannetzky for providing technical material during the establishment of the experimental procedure.

## Author Contributions

IS established the experimental procedure. TT developed the Fiji script. IS and TT developed the Rpackage, designed the figures and wrote the manuscript.

## Supplementary Material

The supplementary material for this article can be found online at:

## Supplementary Material

Video 1 Molting event larval stage one to larval stage two; recorded at 3 fpm, played back at 3 fps

Video 2 Molting event larval stage two to larval stage three; recorded at 3 fpm, played back at 3 fps

**Figure S1** Scheme of the Fiji macro script. Frames were distributed in packages of 1080 frames (equivalent to 6h at 3fpm) and then cropped in regions of interest (ROIs). ROIs were analyzed with specific parameters. Parameters for analyzed particles were saved in a csv-file. Batches at the border of two developmental stages were analyzed with parameters of both respective stages.

**Figure S2** Scheme of the R script. Csv-files were loaded in R and combined into a newly created list. Data were evaluated in two steps. First, fitting settings where selected and then objects where analyzed with these settings. After final evaluation of the data, new csv-files were saved and results were plotted using ggplot2.

**Figure S3** Molting events of individual first instar larvae. Note the phases of low activity in all images. Low activity indicates ecdysis behavioral sequence of *D. melanogaster*.

**Figure S4** Molting events of individual second instar larvae. Note the phases of low activity in all images. Low activity indicates ecdysis behavioral sequence of *D. melanogaster*.

