## Supplementary figures and images for "The PEDtracker: an automatic staging approach for *Drosophila melanogaster* larvae"

### FigureS1.tif

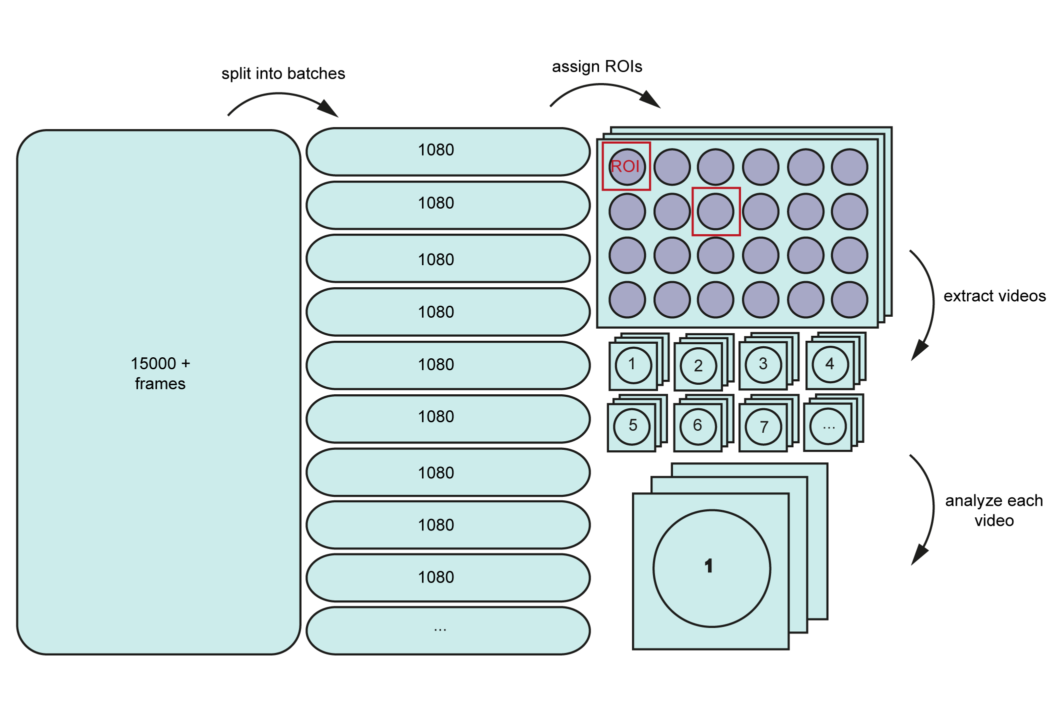

### FigureS2.tif

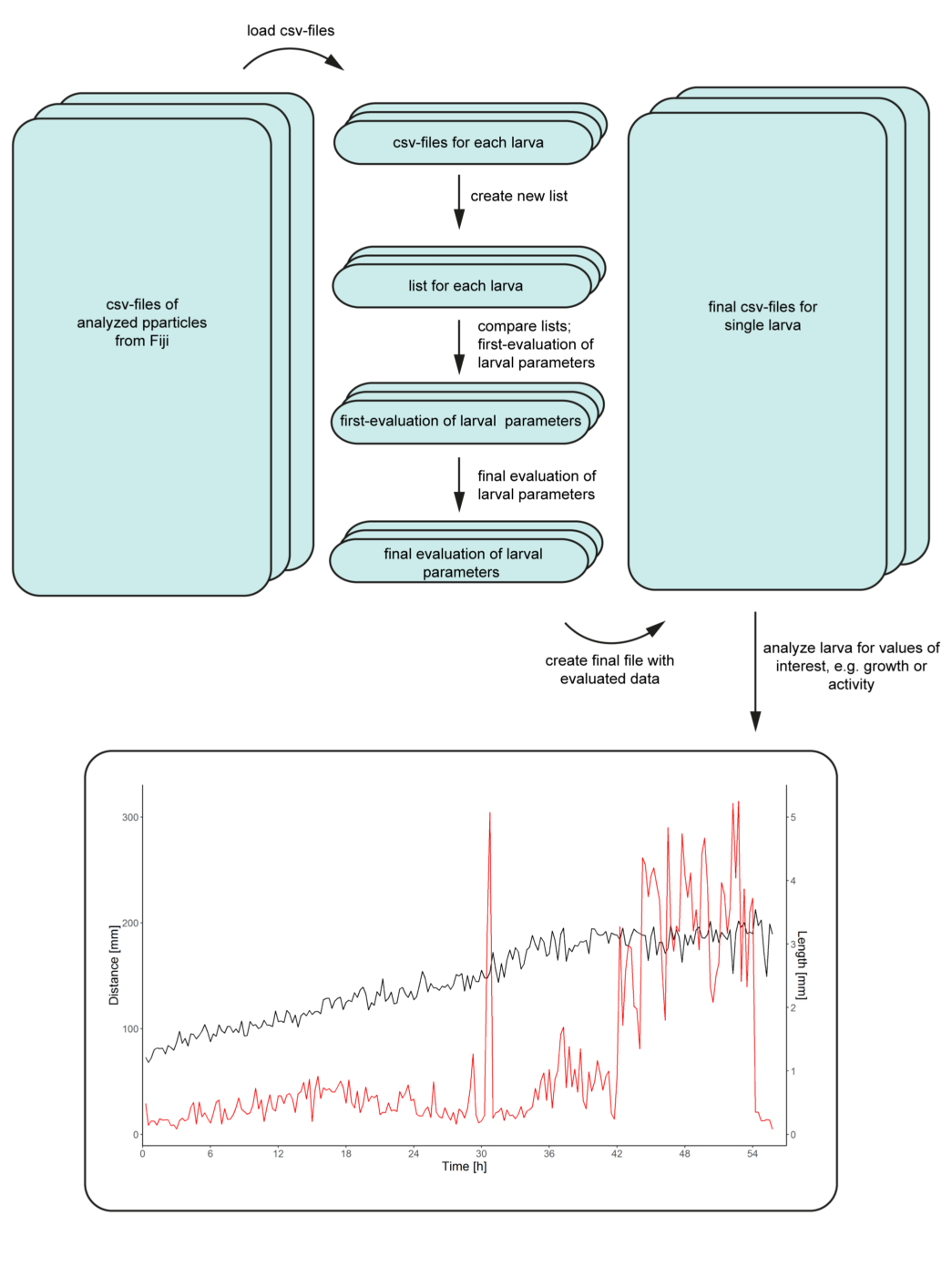

### FigureS3.tif

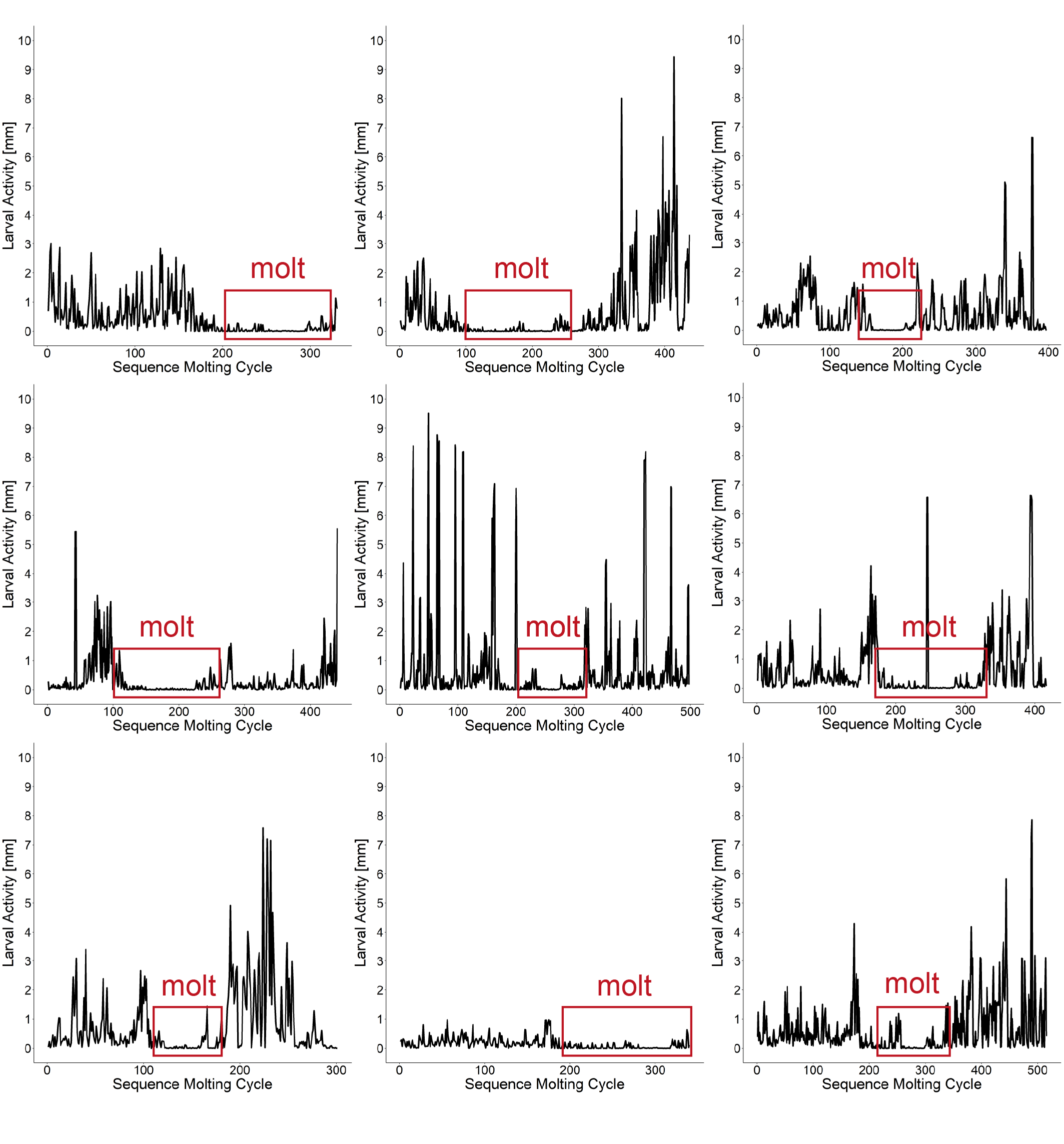

### FigureS4.tif

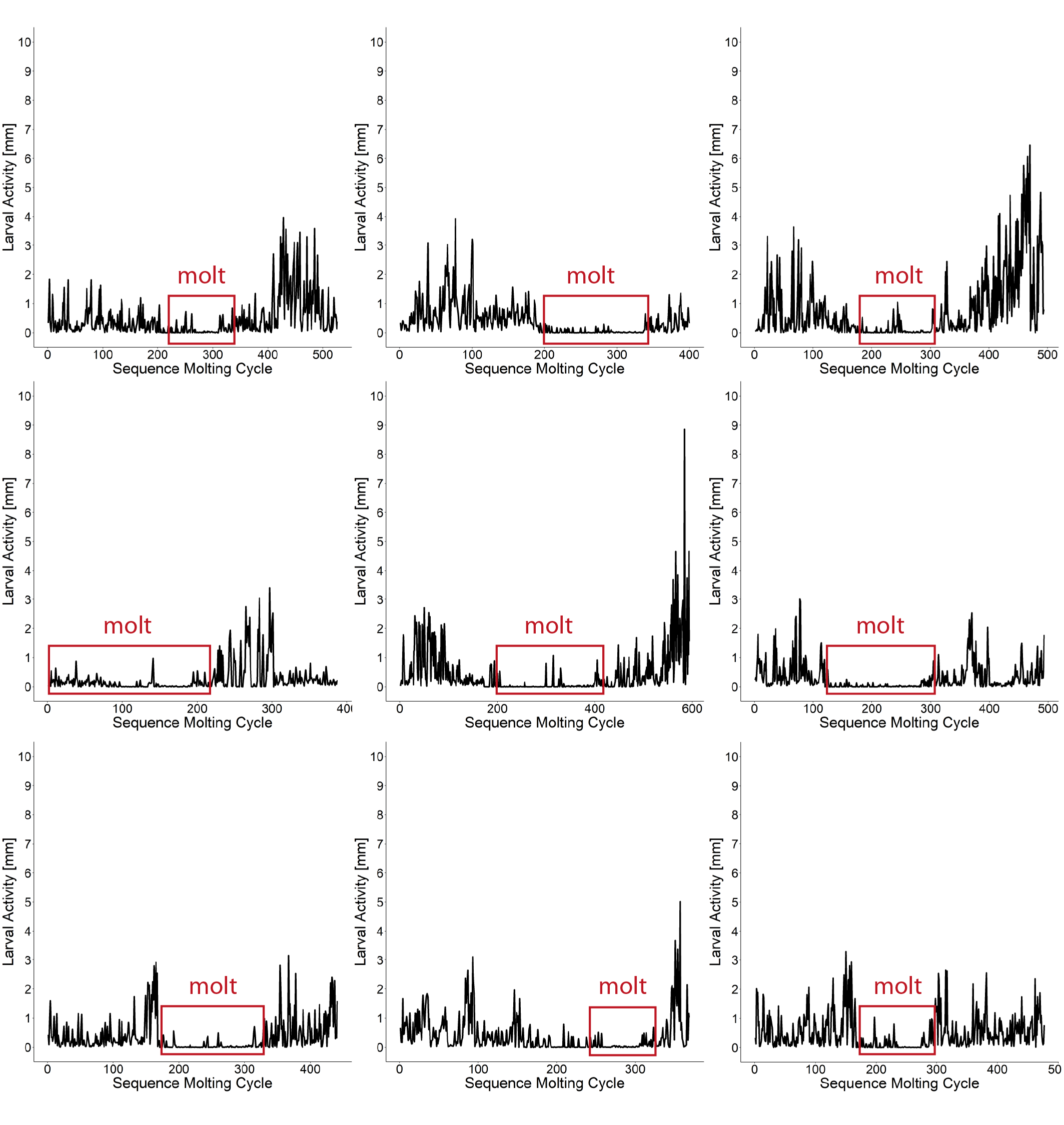
